# Isolating selective from non-selective forces using site frequency ratios

**DOI:** 10.1101/2024.09.13.612810

**Authors:** Jody Hey, Vitor A. C. Pavinato

## Abstract

A new method is introduced for estimating the distribution of mutation fitness effects using site frequency spectra. Unlike previous methods, which make assumptions about non-selective factors, or that try to incorporate such factors into the underlying model, this new method mostly avoids non-selective effects by working with the ratios of counts of selected sites to neutral sites. An expression for the likelihood of a set of selected/neutral ratios is found by treating the ratio of two Poisson random variables as the ratio of two gaussian random variables. This approach also avoids the need to estimate the relative mutation rates of selected and neutral sites. Simulations over a wide range of demographic models, with linked selection effects show that the new SF-Ratios method performs well for statistical tests of selection, and it performs well for estimating the distribution of selection effects. Applications to two populations of *Drosophila melanogaster* reveal clear but very weak selection on synonymous sites. For nonsynonymous sites, selection was estimated to be far weaker than previous estimates for *Drosophila* populations.

## Introduction

Population genomics is often a science of sifting signal from noise as investigators regularly seek to distill the signs of natural selection from the confusing patterns of variation that arises from other factors (1-4). These other factors are quite diverse with some being especially noise-like (genetic drift, recombination and gene-conversion events, and the effects of linked random mutations) and others that have a directional (i.e. non-random) component, as occurs when the demographic history departs from the assumptions of the investigator’s model. Here we describe a new approach for estimating the distribution of selection coefficients acting on mutations, but that does so while largely sidestepping the confounding effects of all these other factors.

For questions about selective effects investigators have often employed a classic body of theory on the distribution of allele frequencies (5, 6), also known as the site frequency spectrum, or SFS. An important theoretical advance was the realization that the count of observed sites with an allele at a particular frequency could be modelled as a Poisson random variable, with an expected value that depends on a particular model of directional selection and population demography (7, 8). These Poisson random field (PRF) models provide accessible likelihood formulae, not only for the estimation of single selection coefficients, but also for the estimation of the distribution of selection coefficients (9-11).

The original PRF work was limited to constant size Wright-Fisher (WF) populations. To allow for departures from WF models, these methods have been adapted for joint estimates of selection and demography under models of population size change (12-15) and with gene flow between subpopulations (16). Nevertheless, real populations can have histories that vary in many ways not accounted for with these methods. Nor do such methods account for the many other non-selective non-demographic factors that can shape the distribution of allele frequencies, including mutational biases and gene conversion biases, variation in mutation and recombination rates across the genome, as well as linked selective effects such as background selection and selective sweeps. Finally, the structure of the sample across subpopulations can have a very large effect on the site frequency spectrum, one that may easily not be appreciated if there are unknown subdivisions within the sampled population(s).

One way to improve upon methods that attempt to jointly estimate selection and other factors is to include in the analysis a set of neutral control variants that are thought to be affected only by non-selective factors. In particular, Eyre-Walker and colleagues developed a method that uses the joint likelihood of selected and neutral variants, and represents shared, non-selective factors by a series of nuisance parameters, one for each frequency bin (10). This approach has been adopted in a number of studies and applications (17-20).

However the method of using both selective and neutral sites, while potentially solving one problem, introduces another complication, which is that the respective mutation rates for each of the two classes must be estimated. This can be done using a previously estimated mutation rate and by assuming a particular demographic history, or by jointly estimating that history. However in most applications it is handled by including in the likelihood function factors *L*_*N*_ and *L*_*S*_, the number of sampled sites where a neutral or selected mutation could occur, respectively. If these values are known without appreciable error, then the counts of sites that are invariant with respect to a sister species can be obtained, and these can be used to estimate the two mutation rates. One important benefit of this approach is that the divergence measures can be used in turn to estimate the rate of adaptive substitution (9, 19-21).

For methods that do not include neutral controls or divergence between sister species, and rely only upon the frequency distribution of polymorphisms, the actual mutation rate is not a parameter of much interest. In these cases, it is the changing relative height of the polymorphism count across frequency bins that informs on the effects of selection. However, the methods that use both selected and neutral variants all depend on knowing or successfully estimating the underlying mutation rate, which often depends on knowing the value of the number of sampled neutral positions, *L*_*N*_. This value will typically include a large number of invariant positions. However if a subset of these is not variable because they are actually under selective constraint, then *L*_*N*_ value will be too large. Another issue that arises when using *L* values as a means to include divergence in the analyses, is that non-selective factors may have changed over the course of the divergence process (10).

Here we take a new approach to using a neutral control set for isolating the effects of direct selection on a set of variants. But unlike other methods that depend on estimates of the overall mutation rates to selective and neutral variants, our method does not depend on estimates of the mutation rates for each class of variant. The method depends not at all on estimates of species divergence or estimates of the number of sampled selected and neutral positions.

## Results

### Model

We consider both a neutral set and candidate selected set of biallelic single-nucleotide polymorphisms (SNPs) sampled from *n* genomes. Assume for the moment that the derived allele at each SNP is known, such that each SNP can be represented simply by the number of derived alleles that occur in the sample, a value that varies between 1 and *n* − 1. Under a Poisson random field model, the focus is on the expected numbers of SNPs at each possible sample frequency. For both sets of SNPs the expected counts with *i* derived allele copies are the product of two terms: the first of these accounts for the rate of incoming mutations, denoted as *θ*/2 and *θ*_*s*_/2 for neutral and selected respectively; the second term, denoted as *𝒩*_*i*_ and *ℱ*_*i*_ respectively, are both functions of non-selective effects (i.e. population demography, linked selection effects, etc.) and in the case of *ℱ*_*i*_, the direct effects of selection. Thus, the number of neutral SNPs in bin *i* follows a Poisson distribution with expecation *𝒩*_*i*_ *θ*/2, while for selected SNPs the expectation is *F*_*i*_ *θ*_*s*_/2. For simple diploid Wright-Fisher populations, with free recombination, and lacking any mutational biases, the terms for incoming mutations, *θ*/2 and *θ*_*s*_/2, as well as *𝒩*_*i*_ and *ℱ*_*i*_, are well understood and can be specified as functions of a small set of parameters. Specifically: *θ* = 4*N*_*u*_ where *N* is the population size and *u* is the neutral mutation rate; *θ*_*s*_ = 4*Nu*_*s*_, where *N*_*s*_ is the mutation rate to selected alleles; and *𝒩*_*i*_ = 1/*i*. The quantity *ℱ*_*i*_ is a function of the population selection coefficient of the derived allele, *γ* = 2*Ns*, and is found by integration of a term for the distribution of selected allele frequencies in the population over the probability of sampling *i* alleles in a sample of size *n* (7):

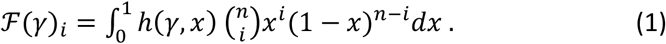

Where *h*(*γ, x*) is the expected density for derived alleles at frequency *x* in the population:

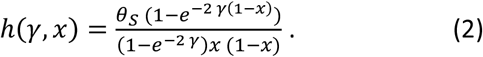

The approach can readily be extended to accommodate a distribution of fitness effects (DFE). Following Boyko et al., (9) if we have a probability density Pr(*γ* = 2*N*) = *g*(*γ, ϕ*), where *ϕ* contains the parameters for the DFE, then an SFS generated under selection will have an expected count for *i* sampled alleles of *ℱg,i θ*_*s*_/2, where

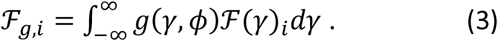

We would like to adapt these methods to problems well outside of the constraints of Wright-Fisher assumptions, to include models with complex histories, and do so without additional parameters for demography or any other non-selective factors that might shape the SFS.

For a data set of unknown history, the expected counts can still be envisioned as the product of a term for incoming mutations and a term for the fraction that are sampled in bin *i*. Now let us suppose that whatever the history, that the ways that non-selective effects shape the expectation for bin *i* can be mostly captured in a term *a*_*i*_, and that this term applies to both neutral and selected SNPs. Further suppose that the effects of selection can primarily be captured separately from the effects captured in *a*_*i*_, by redefining *ℱ*_*g,i*_ as a function not strictly of the probability density of *γ* = 2*Ns*, but rather by shifting the meaning of *γ* so that it is a function, not of census size, but of effective population size (i.e. *γ* = 2*N*_*e*_*s*). We can use *N*_*e*_ as well in redefining the terms for the incoming mutations, *θ* = 4*N*_*e*_*u* and *θ*_*s*_ = 4*N*_*e*_*u*_*s*_. Then for our population of unknown history, the expected count of neutral mutations in bin *i* will be *a*_*i*_ *𝒩*_*i*_ *θ*/2 and for selected mutations it will be *a*_*i*_*F*_*g,i*_ *θ*_*s*_/2.

The motivation for supposing that the bulk of non-selective effects can be separated from the effects of selection is that it opens the door to using the ratio of counts. Of course, the reality is that selection does interact with demography and other factors to shape the distribution of allele frequencies, but a ratio-based approach may still be a useful approximation. Then, for a model of arbitrary non-selective factors, for which the departures from a simple WF model can be capture in a term *a*_*i*_, and by using effective population size rather than census size, the ratio of expected counts in bin *i* is:

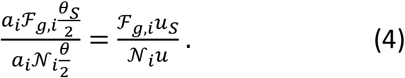

In other words, by taking the ratio of counts, we may be able to work with the standard WF-PRF theory to estimate selection parameters while ignoring whatever non-selective effects might also have shaped the site frequency distribution.

Let *z*_*i*_ be the observed ratio of selected to neutral counts in bin *i*. If we knew the probability of *z*_*i*_ as a function of the distribution of the incoming mutation rates and the DFE, *p*(*z*_*i*_, *ϕ, θ*_*s*_, *θ*), then we could estimate the several unknowns (i.e. *θ*_*s*_, *θ* and *ϕ*) by maximizing the log-likelihood:

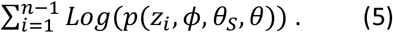

The use of a ratio of candidate selected counts to neutral counts is also supported by the fact that the absolute number of SNPs, and thus the components of *θ* and *θ*_*s*_ corresponding to the rate of incomings mutations, can largely be set by the investigator when working with whole genome data. In other words, in many contexts, *θ* and *θ*_*s*_ are, individually, nuisance parameters. In fact, it turns out that it is practical to work without attempting to estimate both mutation rates, but instead to estimate just the ratio of mutation rates.

### The probability of an observed ratio, *z*_*i*_

Let the data be taken as a set of *n* − 1 ratios, where *z*_*i*_ is the ratio of the observed count of selected SNPs in bin *i* to the corresponding count for neutral SNPs. If we use the basic WF-PRF model for both the numerator and denominator, as in (1), then *z*_*i*_ will be the ratio of two Poisson random variables.

However, as these are each discrete random variables, their ratio is also a discrete random variable with a complicated probability density (22). For a more tractable likelihood calculation, we assume that both the numerator and denominator counts follow a gaussian distribution in which the expectations and variances are both equal to the expectations of their respective Poisson distributions. Clearly the gaussian is continuous, but otherwise a gaussian with expectation and variance *X* provides quite a good fit for a Poisson density with parameter *X* for values even as low as 10. With genome data it is often possible to have more than 10 SNPs in each bin, even for selected SNPs and even for high values of *i*.

For the probability density of the ratio of two normal distributions, we use the formulation by Díaz-Francés and Rubio (23). Because the expectation equals the variance in the case of a Poisson random variable, and thus also for the gaussian distributions used in the approximation, this formulation becomes:

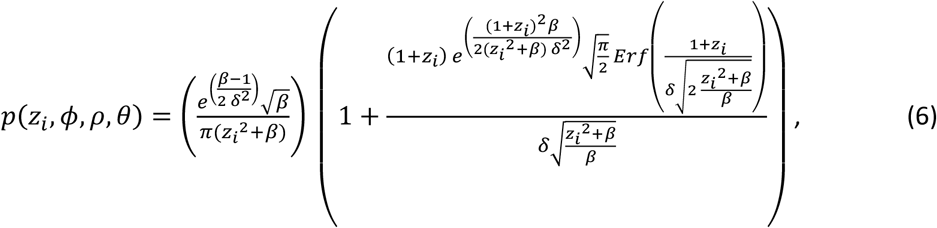

where 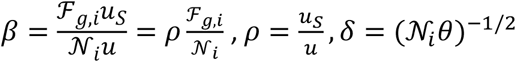, and *ErF()*is the error function. The only place that a mutation rate term appears outside of a ratio is in *δ*, which shapes the variance of the density but has modest effect on the expectation. As the individual mutation rate terms are nuisance parameters, we can consider integrating over *θ* to remove the *δ* term and generate a simpler version of (6) that is a function of only the DFE and *ρ*. This we do, numerically, over two orders of magnitude of values of *θ* that is centered on a log scale at Watterson’s estimate of *θ*, leaving us with a density that is a function of the data, the ratio of mutation rates, and the parameters of the DFE: 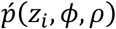. Then the log-likelihood of a set of ratios for unfolded SFSs is simply

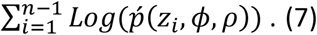

For the folded distribution we substitute 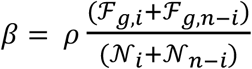, and the log-likelihood is summed over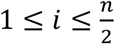. With large amounts of data, expression (7) should open the door to all the applications for which likelihoods are suitable, including parameter estimation, hypothesis testing and model choice. We have named the use of expression (7) for estimating *ρ* and *ϕ* as the “Site Frequency Ratios” method, which we abbreviate as SF-Ratios.

### Working with the ratio of mutation rates, *ρ*

A key parameter is *ρ*, the ratio of the total rate of mutation to selected alleles over that portion of the genome screened for selected variants, divided by the corresponding rate for neutral variants. This ratio should not depend on any factors other than these mutation rates (i.e. no effects of selection, demography, linkage, or sampling geography). However, *ρ* will depend on the relative sampling effort of the two classes of SNPs. For example, if SNPs in the selected pool were sampled from a smaller fraction of the genome than for neutral alleles, then we would expect an estimate, 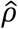, to be below one.

Despite being a function of sampling effort, an estimate of *ρ* can be useful in a at least two different ways. One use of 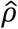 is to consider it together with the total counts of selected and neutral SNPs, which we denote by *X* and *Y* respectively. If in fact both sets were strictly neutral, then a useful estimate would simply be 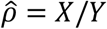. But when there is selection on the SNPs in the numerator, the difference between *ρ* and *X*/*Y* is caused by a difference in rates at which mutations that occurred in the population where unsampled when the data were collected. In particular, deleterious mutations will have a higher chance of going unsampled, on average, relative to neutral mutations, because they have been lost from the population or are more likely to be at extreme allele frequencies. Let *λ* be the probability of not sampling a selected mutation, relative to the probability of not sampling a neutral mutation. Then *ρ* = *λ X*/*Y* and we can estimate the relative probability of sampling as

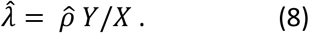

For example, suppose that *X*/*Y* = 0.2 and 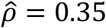, then 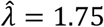, which tells us that a selected mutation is about 75% more likely to go unsampled, relative to a neutral mutation. *λ* can be considered as a measure of the variation that is missing due to the direct effects of selection on deleterious or beneficial alleles. Missing variants will include those that have been lost or fixed, as well as those that fall in frequency ranges that are less likely to be sampled from.

Another use of 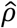 is to apply it to the estimation of selection parameters for other populations of the same or related species. To see this, suppose that we have a value of 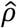 for one population for which we are relatively confidant that the circumstances are favorable for an accurate estimate. And now allow that there is a second population of the same or closely related species for which we have SNP counts based on the same sampling process as the first population, but for which we are more doubtful that the approach of using the ratios of counts will work for estimating selection parameters. Because the two populations are closely related, and SNPs have been sampled from the same genomic regions, we can assume that the true value of *ρ* is the same for both populations. Then we can take 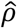 for the first population, and fix *ρ* for the second population at that value, with the hope that the estimates of selection parameters for the second population will be more accurate than if *ρ* were also being estimated for that population.

## Simulation Results

### Qualitative Assessment of SF-Ratios

Simulations were conducted to qualitatively assess the underlying rationale of using the ratio of selected to neutral SFS values to isolate the effects of selection. We simulated SFSs, and the selected/neutral SFS ratios, for several demographic models, and compared them to the basic WF model of constant population size. Figure 1A shows mean values of simulated folded SFSs under a constant WF model and three demographic models, for both neutral mutations and mutations with selection coefficients drawn from a lognormal distribution. To aid comparison among models, which vary in their absolute numbers of polymorphic sites, for each model, the values shown are scaled relative to the count for the singleton bin (i.e. *i* = 1, just one observed copy of the rare allele) for the corresponding neutral model, and then plotted on a logarithmic scale. For this demonstration the ratio of the rate of incoming mutations was set to 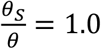. Comparison reveals how the selected sites have an SFS that departs greatly from the neutral case, regardless of the model, and it shows how the models vary considerably in their SFSs, both with and without selection. However, the ratios of these same values, of selected to neutral SFSs, are quite similar for the different demographic models (Fig 1b).

**Figure 1.**
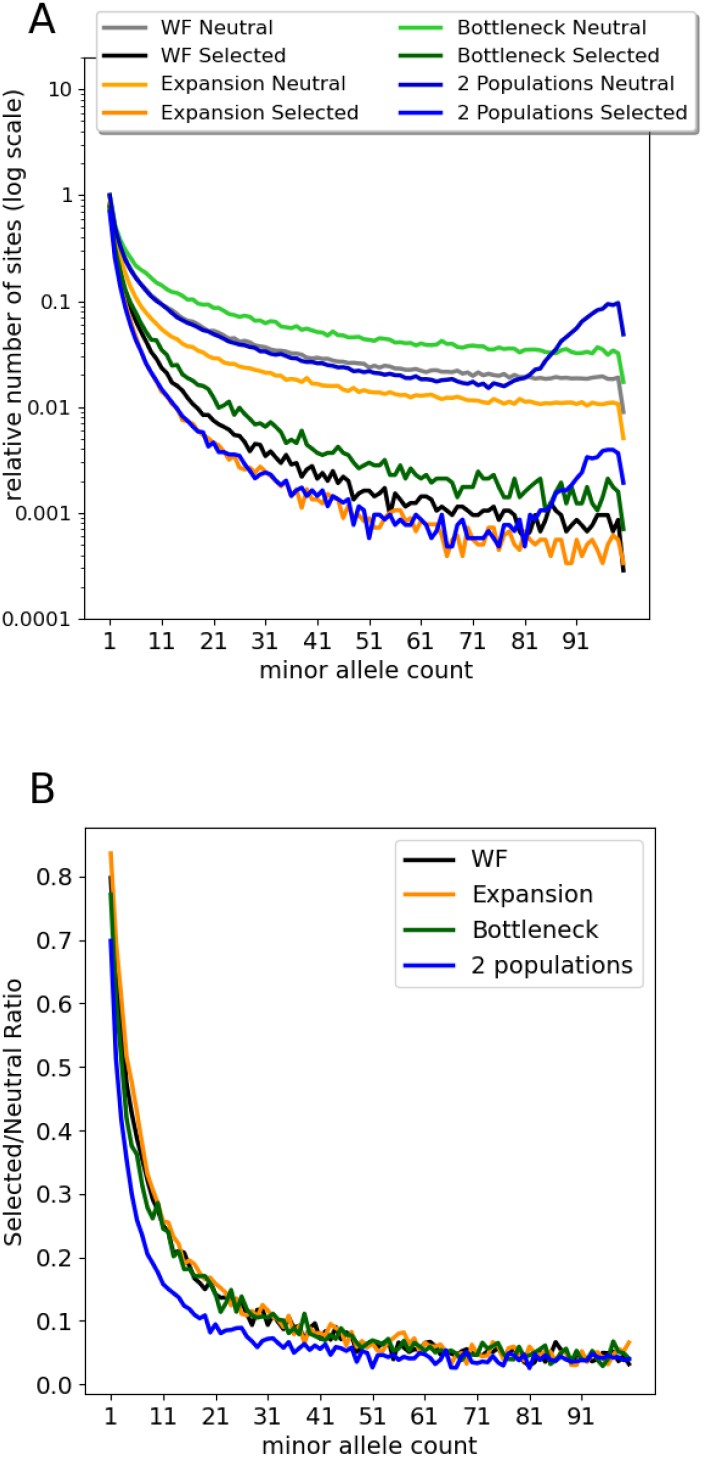
Comparison of SFSs and Selected/Neutral ratio for Wright-Fisher (WF) and other demographic models. Selection model: 2*Ns*∼Lognormal(3.0,1.2) (see methods), with expectation -40.3. For all simulations, 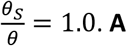. Folded *θ* SFS values for a sample of 200 chromosomes, scaled to the value for allele count 1. **B**. The ratio of selected to neutral counts for folded SFSs.

### Performance on Simple Tests of Selection

One kind of application of PRF theory is to use a likelihood ratio test to determine whether a WF model that includes selection provides a significantly better fit to a data set than a strictly neutral WF model (8). Such tests can be quite powerful, however, without a way to control for the non-selection-based effects on the SFS, they will be sensitive to virtually any departure from a WF model.

We considered whether the likelihood in expression (7) can also be used for simple tests of selection by comparing the performance to that based on regular SF-based likelihoods. Computer simulation of SFSs with selection were examined using likelihood ratio tests and subjected to power analyses and receiver operator characteristic (ROC) curve analyses. The same kinds of analyses were then conducted with each simulated data set transformed into a series of ratios using a simulated neutral control and then examined using a likelihood ratio test.

Figure 2 shows results for the statistical performance of basic tests of the hypothesis that the sampled data came from a population with *γ* = 0. Each panel compares simple WF-PRF simulations in which the data were sampled from a Poisson distribution for a sample size of 100 genomes, with *θ*_*s*_ = 500, to the SF-Ratios method in which each data set of selected alleles is paired with a neutral data set of 100 genomes and with *θ*_*s*_ = *θ* = 500. In each case, statistical significance was determined by comparing the 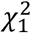 value for each of three false positive rates (0.05, 0.01, 0.001) to twice the log of the likelihood ratio. These are

**Figure 2.**
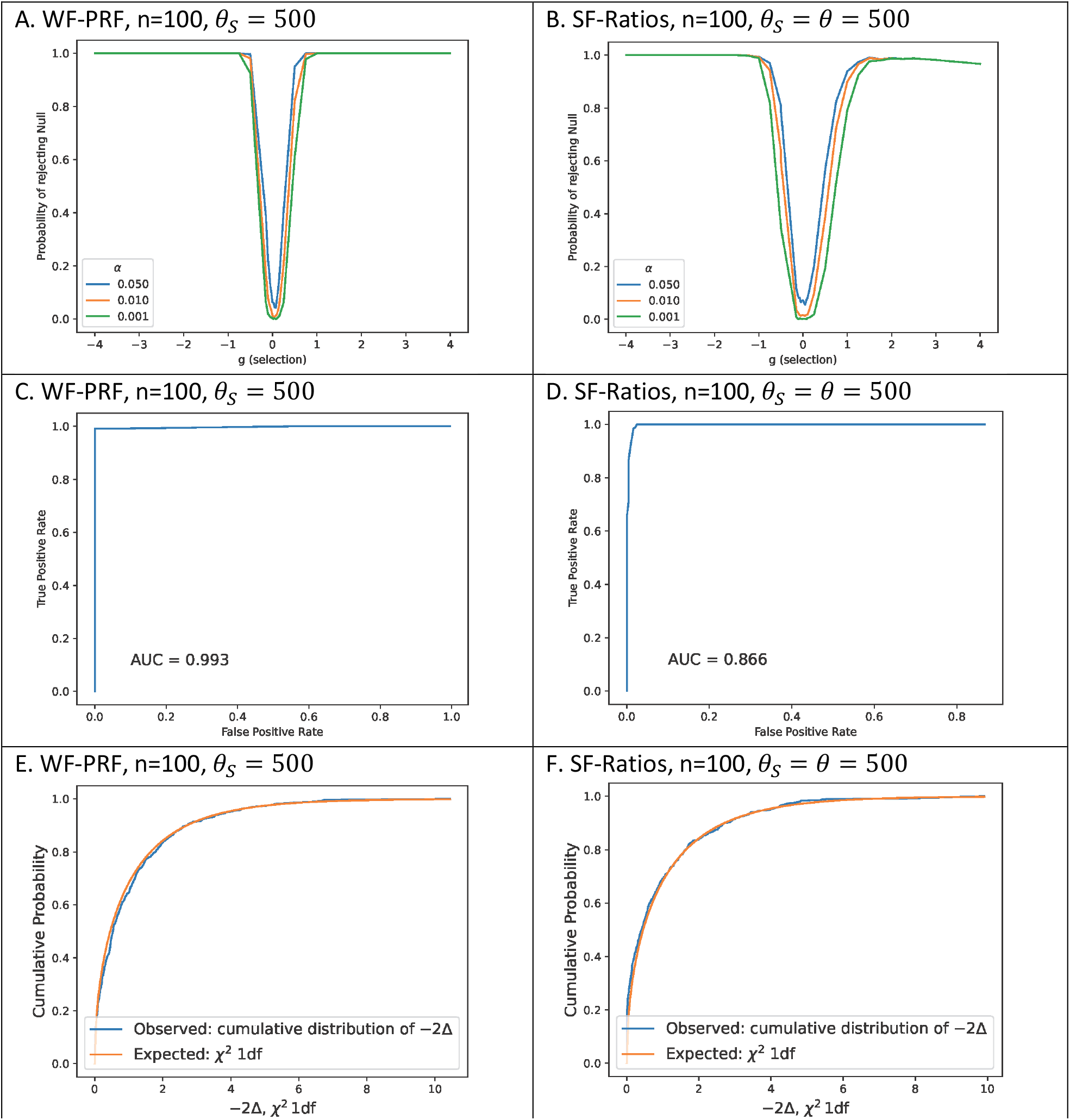
Top row (panels A, B). The probability of rejecting the null (neutral) when the alternative model (selected) is true for different probabilities of false positive (α) and varying strengths of 2Ns. Middle row (panels C, D). Receiver operator characteristic (ROC) curves, with area under the curve (AUC). Bottom row (panels E, F). Cumulative observed distributions of the likelihood ratio test statistic, with χ^2,1df comparison for sets of 500 simulations.

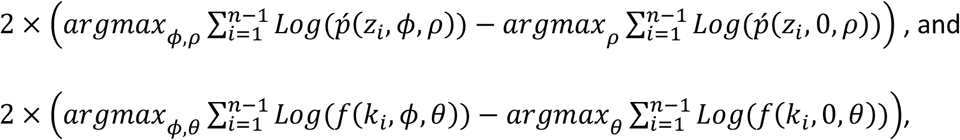

Where 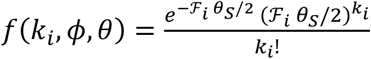, for SF-Ratios and WF-PRF, respectively.

Panel A of figure 2 shows the statistical power for −20 ≤ *ϕ* ≤ 20. Panel B shows ROC curves for SF-Ratios and WF-PRF for the same hypotheses and test distribution 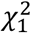 as shown in panel A. For this analysis half of the simulations (500) were done when the null hypothesis is true (*ϕ* = 0) and half when −100 ≤ *ϕ* ≤ 1, with values sampled uniformly at random. Panel C compares the observed distribution of the test statistic to the cumulative 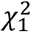 distribution for 1000 simulations when the null hypothesis is true (*ϕ* = 0). Results for smaller data sets, including simulations with low variation and small sample size (*θ* = 50, *n* = 20) and low variation and large sample size (*θ* = 50, *n* = 100), are given in Figures S1, S2 and S3.

In general, the analyses based on the WF-PRF likelihood showed more statistical power, a closer fit of the likelihood ratio test statistic to the expected *χ*^2^ distribution, and higher area under the curve (AUC) in the ROC analyses, compared to the SF-Ratios likelihood tests. However, statistical power was high for both methods for all but the smallest selection coefficients; AUC was high for both methods for all 3 sampling schemes, and the *χ*^2^ approximation fitted well except for the smallest sample sizes, even when the fit was accurate in the tail of the distribution associated with the smaller false positive rates. However, even though the SF-Ratios likelihoods performed well, it is important to recognize that they are all based on twice as much data as the corresponding SF-based likelihoods, in that, each SF-Ratios data set had both a selected SFS and a neutral SFS.

### Estimator Bias under Wright-Fisher and other Demographies

Forward population genetic simulations were conducted to include linked neutral and selected mutations under several non-WF models. For fixed values of *γ*, simulations were conducted under a constant population, recent population expansion, recent population bottleneck, and population structure. For simulations under lognormal DFE, these same demographic models were used, as well as an African-Origin model of human demographic history (24). Analyses were performed for the African sample SFSs, and for the SFSs from pooling the three populations (Africans, Europeans, and East-Asians) before sampling of genetic variation used to obtain the SFSs.

Figure 3 shows boxplots of estimates for each of a series of selection values for each model. In general, weak selection (beneficial or deleterious) was estimated well in all the models, whereas positive 2*Ns* ≥ 10 was underestimated in all models (i.e. estimate values were closer to zero). Strong deleterious selection was overestimated (i.e. estimated values were more negative) in the bottleneck and population structure models.

**Figure 3.**
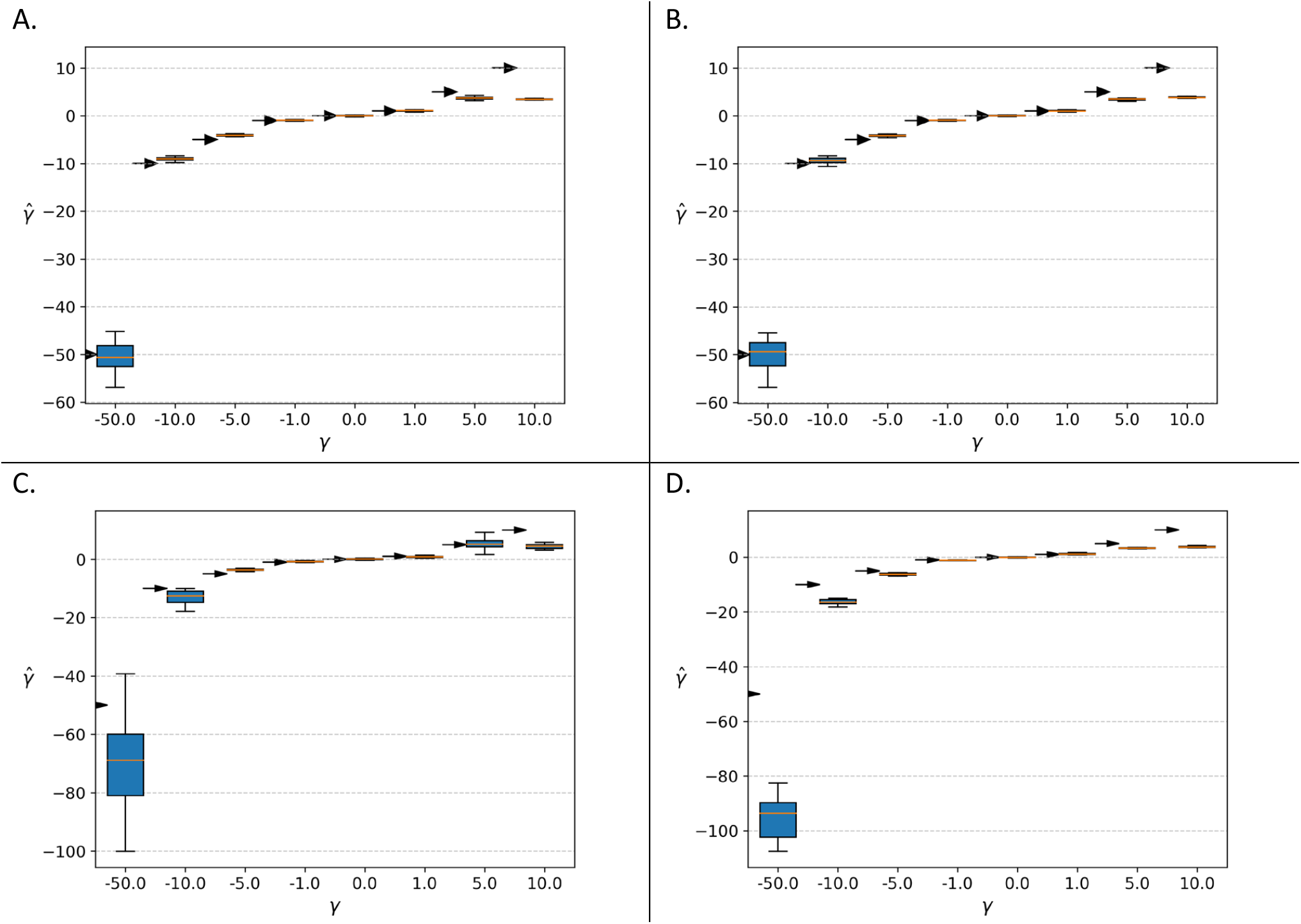
Boxplots of *γ* values estimated for simulated data generated under different demographic models. For each box, an arrow indicates the location of the true value. A. Population with a constant size. B. Population expansion. C. Population bottleneck. D. Two divergent subpopulations.

Figures 4 and 5 shows results for simulations under a lognormal DFE, each extending from highly negative values to +1, in order accommodate strictly neutral mutations, some beneficial mutations, and as wide a range as necessary of deleterious mutations. The lognormal distribution is parameterized using the expected value, *μ*, and the standard deviation, *σ*, of the random variable’s natural logarithm. We considered four sets of parameter pairs, each of which generates a curve with a peak near zero, but with increasingly negative mean values (Fig 4 A). Estimator bias for *μ* and *σ* is shown in 2D boxplots (Figs 4B thru 4G) and for *ρ* in Figure 5.

**Figure 4.**
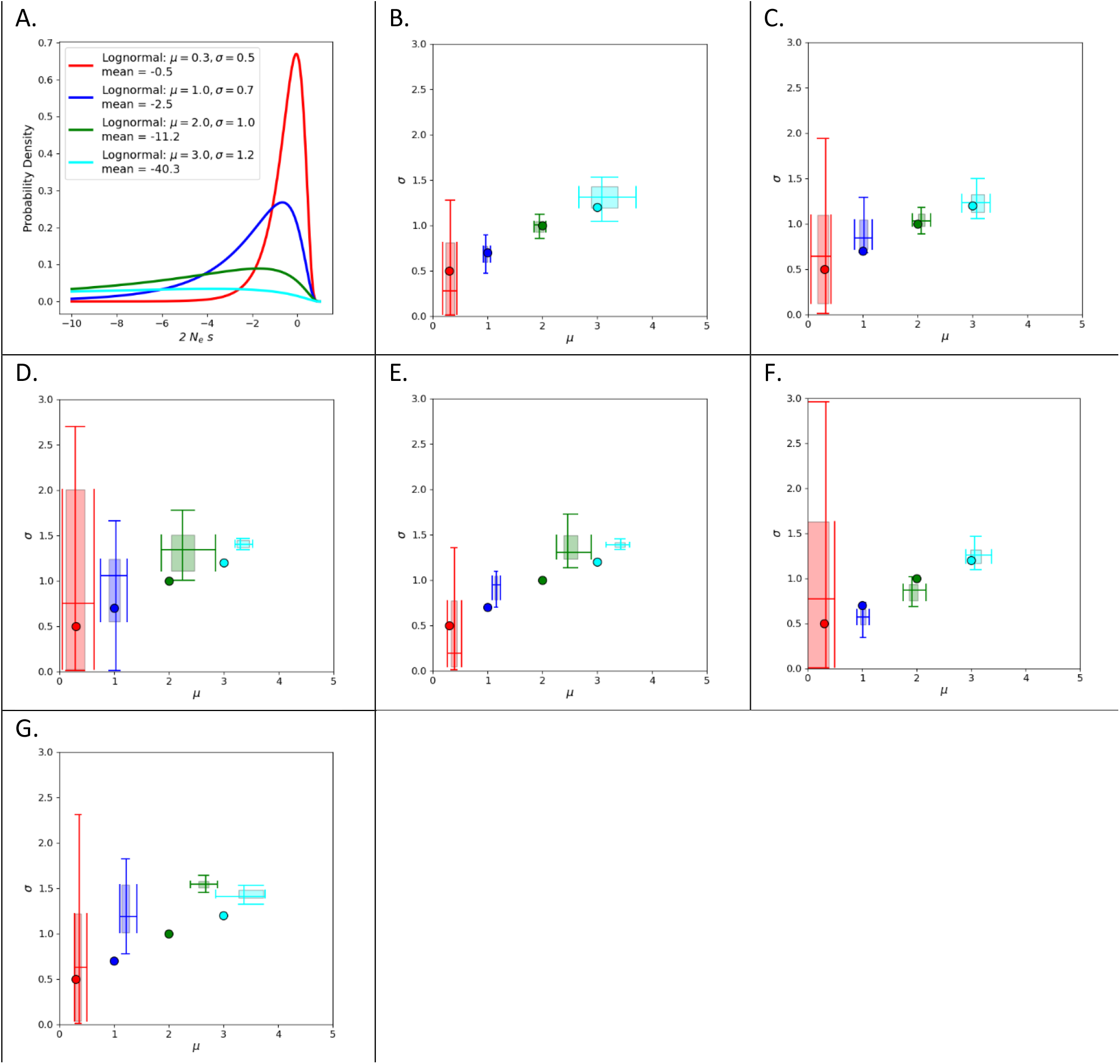
Estimator performance for lognormal parameters. For each simulated data set, *γ* values were drawn from one of 4 lognormal distributions. 20 data sets were simulated for each demographic model and each *γ* distribution. A. lognormal distributions, for random variable *x*, (0 < *x* < ∞), 2*N*= 1 − *x*. B. Constant population size Wright-Fisher. C. Population expansion. D. Population bottleneck. E. Two populations. F. African sample from an African-Origin model. G. African Origin model, pooled sample.

**Figure 5.**
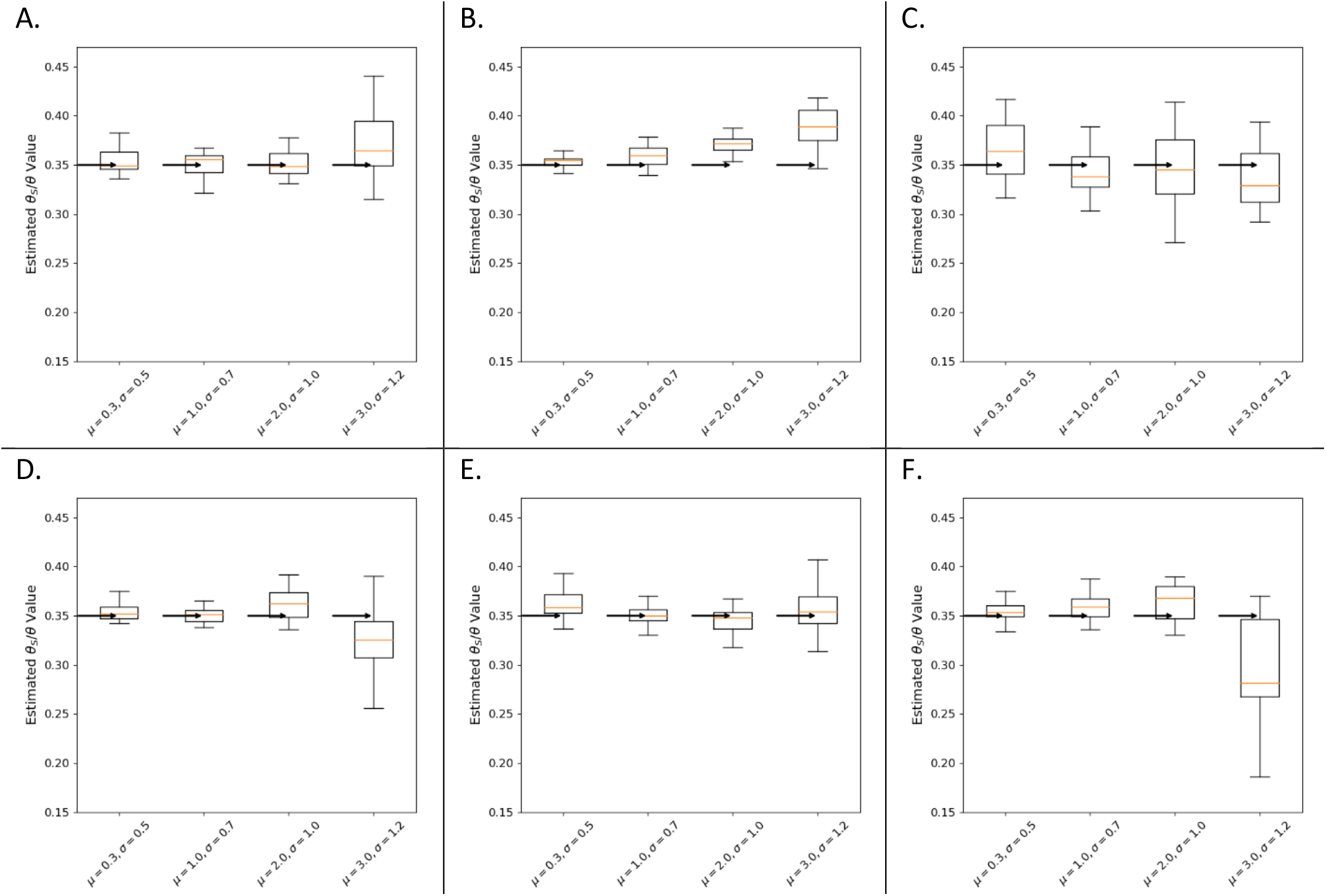
Estimator performance for *ρ*. For each simulated data set, 2*NN* values were drawn from one of 4 lognormal distributions (see Fig. 4). 20 data sets were simulated for each demographic model and each 2*Ns* distribution, each with a true mutation rate ratio of 0.35. A. Constant Wright-Fisher B. Population expansion. C. Population Bottleneck. D. Two populations. E. African sample from an African-Origin model. F. African Origin model, pooled sample.

For the ratio of mutation rates, *ρ*, bias was near zero or small for all models and parameter sets (Fig 5). For the parameters of the 2*Ns* distribution (Fig. 4), statistical bias is low to modest when selection is weak, regardless of the demographic model being used. However for models with strong selection, the statistical bias was higher for some demographies, particularly for the bottleneck and African Origin models. The direction of bias for stronger selection tended to place joint parameter estimates further away from the origin in the plane of values (Figure 5), corresponding to estimated distributions that were stronger (more negative mean fitness values) relative to that for the true distributions. Overall, estimates of *ρ, μ* and *σ* tended to be close to the true values for a wide range of selection models for a wide range of demographies, and this is especially true for weaker selection.

### Drosophila Populations

For two population genomic datasets from *D. melanogaster* we generated folded SFSs for both synonymous and non-synonymous SNPs. In each case the neutral control set was generated from short introns (25, 26). To further reduce sources of variance, the neutral control sets were built to reduce variation due to the local sequence context of each SNP. SNPs for the neutral control sets were chosen by identifying for each candidate selected SNP (synonymous or non-synonymous) the closest short intron SNP having matching flanking bases (17).

Figure 6 shows the folded SFSs for both populations for nonsynonymous, synonymous and short-intron SNPs. The relative intensity of selection can be roughly perceived by the flatness of the distributions, with more strongly selected sites showing a greater drop in counts from low frequency bins to higher frequency bins. The ratios with counts for short intron SNPs in the denominator, are shown in Figure 6.

**Figure 6.**
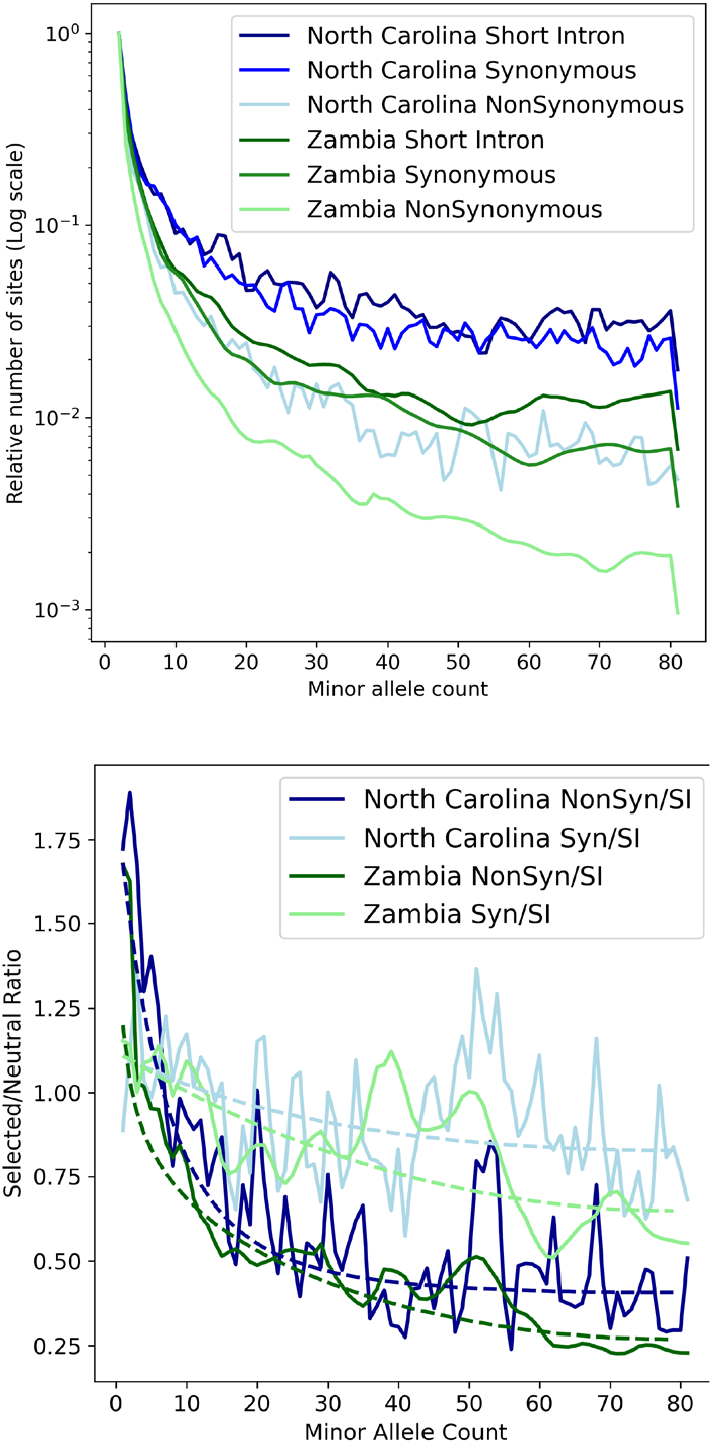
Top. SFSs for two Drosophila populations, each downsampled to n=160 and folded. Values were normalized to that singleton bin count to enable comparisons. Bottom. SFS ratios for both populations, for nonsynonymous and synonymous SFSs divided by the short intron SFS. Expected values generated under the best fit models are shown with dashed lines.

For both populations we fit three types of continuous distributions (normal, lognormal and gamma) as well as each of these DFEs with the addition of a single point mass of up to 50% of the total density. Model fitting involves estimating either three parameters (i.e. *ρ* and density parameters) or five parameters (in the case of an added point mass), in which case the same three parameters are estimate as well as mass location and value. Full results are shown in tables S1 and S2 with the best fitting models shown in Table 1 and Fig. 7.

**Table 1.** Best fitting models.

| Nonsynonymous sites |  |  |  |  |  |  |  |  |  |
| --- | --- | --- | --- | --- | --- | --- | --- | --- | --- |
| Pop | DFE<sup>1</sup> | K<sup>2</sup> | AIC<sup>3</sup> | $\hat{\rho}$ | $\lambda_1$<sup>4</sup> | $\lambda_2$<sup>4</sup> | $p_+$<sup>5</sup> | $\gamma_+$<sup>6</sup> | Mean<sup>7</sup> |
| Zambia | Gamma<br>$\gamma_+, p_+$ | 5 | -1681 | 2.19<br>2.14,2.23 | 4.37<br>3.90,4.85 | 13.40<br>11.75,15.2<br>0 | 0.36<br>0.35,0.37 | -1.60<br>-1.67,-1.52 | -37.2 |
| North Carolina | Gamma | 3 | -1458 | 2.22<br>2.17,2.29 | 0.88<br>0.86,0.90 | 20.3<br>19.1,21.5 |  |  | -16.9 |
| Synonymous sites |  |  |  |  |  |  |  |  |  |
| Zambia | Normal | 3 | -1565 | 1.14<br>1.11,1.16 | -1.01<br>-1.09,-0.93 | 0.63<br>na ,0.98 |  |  | -1.01 |
| North Carolina | Normal | 3 | -1414 | 1.14<br>1.11,1.17 | -1.24<br>-1.43,-1.04 | 3.39<br>2.71,4.32 |  |  | -1.24 |

<sup>1</sup> The best fitting distribution model for 2Ns. If the best model included a point mass, $p_+$ is the value of that mass and $\gamma_+$ is the location (i.e. the 2Ns value) <sup>2</sup> The number of parameters <sup>3</sup> Akaike information criterion (32) <sup>4</sup> depending on which continuous distribution fit best, $\lambda_1$ , $\lambda_2$ are either the lognormal mean and standard deviation, or the normal mean and standard deviation, or the gamma $\alpha, \beta$ . <sup>5</sup> Point mass value <sup>6</sup> Point mass location <sup>7</sup> The expected value of the distribution model for 2Ns

**Figure 7.**
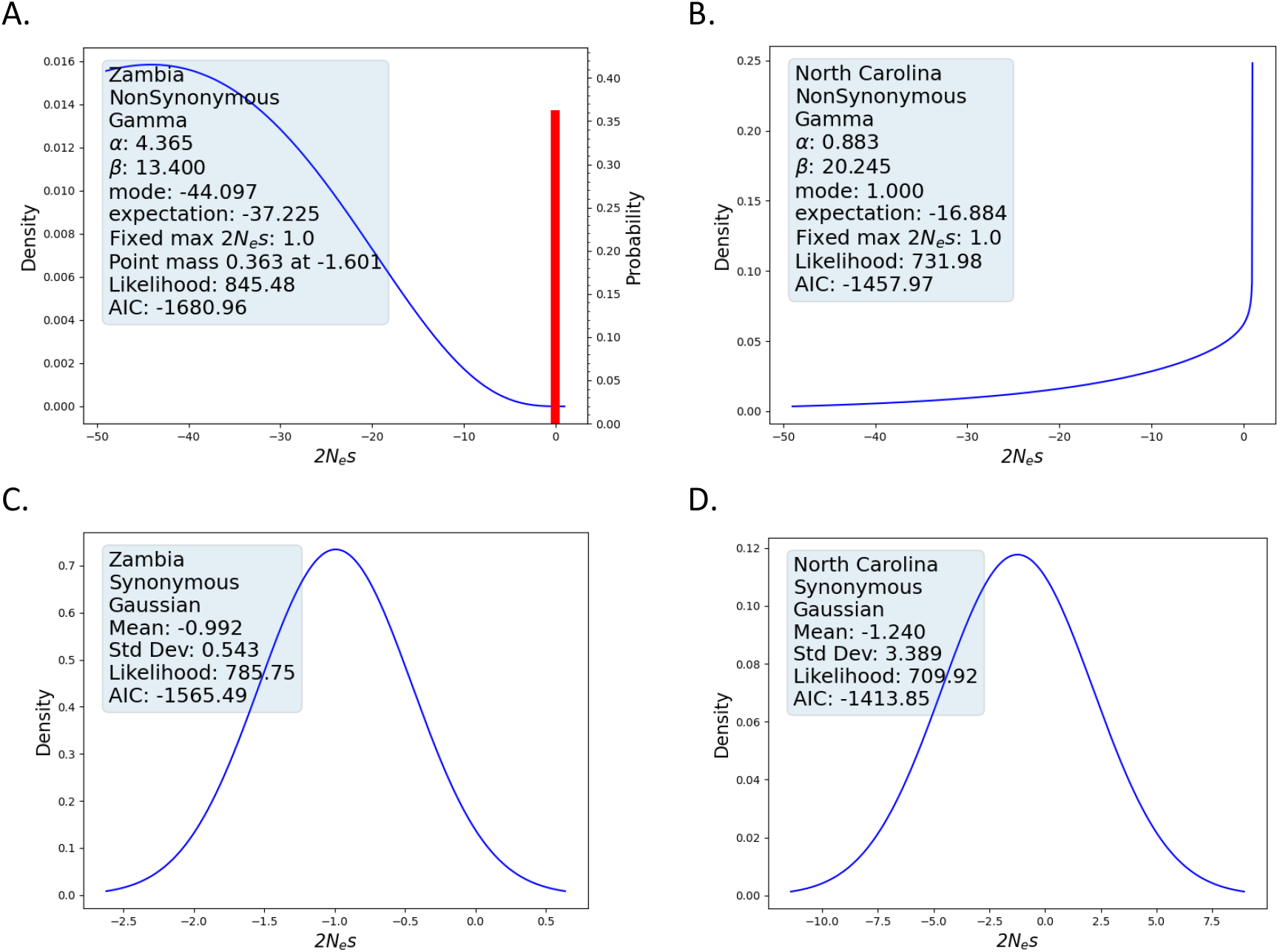
Best-fit estimated 2*Ns* densities for *Drosophila* populations. A. Non-synonymous variation, Zambia. B. Non-synonymous variation, North Carolina. C. Synonymous variation, Zambia. D. Synonymous variation, North Carolina.

The SF-Ratios fitting for nonsynonymous sites, selection appears to have been fairly modest in the Zambia population, with an estimated mean of the *γ* density of -37.2, while that of the North Carolina population is weaker still at - 16.9. Given that the Zambia population has had the larger effective population size, the difference in mean selection strength is in the expected direction.

Supporting these results is the finding that the estimated values of the mutation rate ratio, *ρ*, are quite similar in the two populations (2.19 for Zambia, 2.22 for North Caroline), despite the Zambia population having had a far simpler demographic history than the North Carolina population (27, 28). This value, ∼2.2, is also very close to that expected based on sampling effort, in which non-synonymous changes were sampled over coding portions of exons, while all base positions in short introns where considered (excluding 8 bp on each end). Given this 3 to 1 ratio in sampling effort, we can solve for the fraction, *h*, of coding bases than can mutate to a non-synonymous allele (3 × *h* = 2.2), in which case *h* = 0.7333, which is close to the value of 0.77 that one gets assuming random amino acid and codon usage (29).

The best model for non-synonymous variation in the Zambia population included a large point mass at - 1.6, while the best model for the North Carolina population did not include a point mass, but did have a strong peak at the upper bound. The two distributions appear qualitatively different (Fig. 7), although both share a large density near the upper bound, with an extended tail into negative values.

As described above, we can use 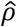 and the counts of selected and neutral SNPs to estimate *λ*, the relative probability that a selected mutation goes unsampled, relative to a neutral mutation. Because we used pairs of intron and exon SNPs, the ratio of the numbers of polymorphic sites (*X*/*Y* in expression (8)) is 1, and so 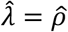 in the present case. For nonsynonymous sites in Zimbabwe and North Carolina the relative number of selected mutations that have gone unsampled, 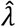, is 2.19 and 2.22, respectively. In other words, nonsynonymous mutations in this population are about 2.2 more likely than neutral mutations to go unsampled because of selection removing them from the population or driving them to extreme allele frequencies.

The SF-Ratios fitting for synonymous sites show a consistent pattern across populations and 2*Ns* densities. Both populations were fit equally well by the Gaussian, lognormal and gamma distributions (Table S2), with the values for the Gaussian density shown in Table 1 and Figure 7. For both populations 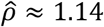 and the mean of the estimated density is near -1, similar to previous estimates (30, 31). Given that the VCF files of both populations were sampled with the same pipeline and protocol, their true values of *ρ* are expected to be very similar as this quantity depends only on the underlying mutation rate and the sampling effort.

## Discussion

The promise of the SF-Ratios method is to enable the estimation of selection intensity without having to consider either divergence between species, or the underlying mutation rates, or the other non-selective factors that will also have shaped polymorphism patterns. Given the simulation results, the method works well across a wide array of demographic scenarios, particularly when selection is not strong. Although not examined here in depth, the approach of using ratios should also be effective for other non-selective factors, in addition to demography, so long as they are shared by the selected and neutral variants. For example, if the two sets of variants are sampled near each other (as in this study), then linked selection effects, including background selection and selective sweeps, will have affected both in similar ways. Similarly, the use of ratios should accommodate factors that affect mutation biases and gene conversion biases if they are shared by selected and neutral variants, as is partly the case in this study. However, if the selected and neutral sets differ because of factors that are not shared, such as differing levels of biased gene conversion due to base composition differences, as occurs in mammals (33), then the ratio-based estimate may suffer.

The SF-Ratios method depends strongly on the quality of the neutral control set of SNPs under three main criteria. The first is selective neutrality, which can be inferred in relative terms using site frequency analyses and divergence patterns (26), but it is difficult to know with certainty. The second criterion is that the neutral set share as many of the non-selective factors as possible with the selected set of SNPs. All parts of the genome share a demographic history, but not all share equally in background selection or mutation and recombination related processes. This second criterion can often be met, at least approximately, by having the neutral SNPs be near to, and interspersed among, the selected SNPs. The third criterion is to avoid increasing the variance of ratios by being sure to record the frequencies of both types of SNPs, selected and control, on the same set of genomes. This criterion arises from the very large effect that unforeseen population structure can have on the SFS of a sample. Consider for example an SFS for 10 genomes sample from a population that, unbeknownst to the investigator, actually includes two divergent subpopulations. In this case the partition of the sample (i.e. the numbers of individuals in each subpopulation) will have a very large effect on the SFS. If the neutral and selected SNPs were drawn from different individuals, then there will be a strong chance that the two sets will have a different partition with respect to the subpopulations, with very large and differing effects on their respective SFSs.

Our study of the Zambia *Drosophila* population can be compared to estimates from other methods that use site frequency data, but that also relied upon mutation rate or divergence measures. Table 2 shows the best fitting model results of three studies, all of which compared lognormal and gamma densities for *γ*. All these estimates suggest much stronger selection (lower 2*Ns* values) than found in our analyses. Huber et al., (34) estimated a mean 2*Ns* of -738.26 in the Zambia sample, while Ragsdale et al., obtained an estimated mean of -2760 on the Zambian sample (35) and another of -7414 on a Rwandan sample (36). All these other studies rely upon estimates of the number of invariant sites as well as fitting of a demographic model. In our method, by using the ratio of two SFSs, one being the SFS of putative neutral sites, we avoid both of these complications.

**Table 2.**
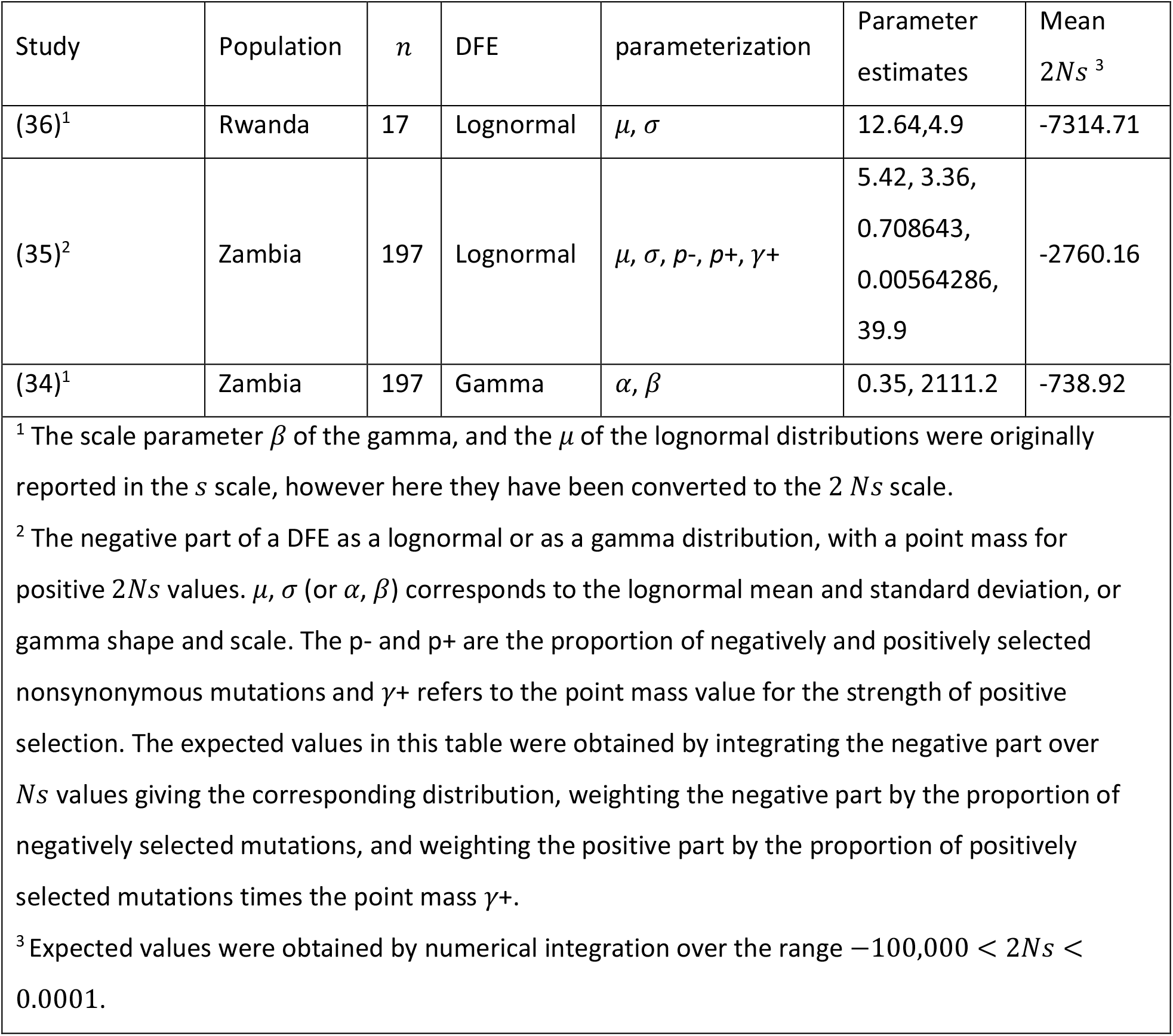
Previous DFE estimates for nonsynonymous mutations.

## Materials and Methods

### Simulations

With SLiM3 (37), we simulated a series of diverse demographies with linked selection effects under both a fixed 2*Ns* and a continuous distribution of 2*Ns* values. For the 2*NN* distribution, we used an inverted lognormal distribution with a maximum set to 1 and a minimum of -100000. For each combination of 2*NN* distribution and demographic model we ran 20 simulations each with a total genome length of 4 megabase pairs (Mbp). To achieve faster running times, we split each genome into 400 fragments of 10 kilobase pairs (Kbp), allowing us to simulate these smaller fragments faster in independent sub-simulations. Each of these 10 Kbp genomic fragments was further divided into eight consecutive neutral/selected pairs of size 810 + 324 = 1134 bp, and a neutral segments of 928 bases. Selected fragments had only non-neutral mutations (i.e. 2*Ns* ≠ 0, while neutral fragments carried only neutral mutations (i.e. 2*N*= 0). Base diploid population size (*N*) was set to 1000, with recombination and mutation rates per base pair of 2.5 × 10^−7^, giving population level rates (4*Nu*) similar to that seen in human populations (i.e. 10^−8^). For each simulation the site frequency spectra (SFS) for all neutral variants sampled from the sub-simulations were summed (as were those for the selected variants), then ratios calculated as the selected count for a frequency bin, divided by the neutral count for the corresponding frequency bin. All simulations began with a burn-in period of 10 *N* generations to allow the population to reach mutation-selection-drift equilibrium (38).

For simulations with changing population size, 2*Ns* values were kept constant by rescaling selection coefficients at each generation in the simulation at which population size changed. We chose this approach, rather than the alternative of keeping *s* constant, to be consistent with our model in which 2*Ns* (or a distribution of 2*Ns*) is fixed. For all models and simulations, we calculated the ratio between each bin of the non-neutral and neutral SFS obtained for a sample n = 200 individuals.

We considered the following demographic models. (1) Constant population size at *N*= 1000. (2). Population expansion, with an initial population of 1000 jumping to 10000, followed by 100 generations before sampling. Population bottleneck, with an initial population of 1000 jumping to 100, followed by 100 generations before sampling. Population structure, with an initial population of 1000, splitting into two populations each of 1000, followed by 5000 generations before sampling equally from both populations.

In addition we simulated the human African-Origin (AO) model as inferred by Gravel et al. (24). This model has some of the features present in the described models, but also some more complex ones. This model includes multiple populations, a bottleneck at the founding of non-African populations, expansion of European and East Asians subpopulations, bottleneck, and population structure as well gene flow among populations. Because the OA simulations with the inferred population sizes are too computationally expensive, we ran a set of neutral simulations to identify the best scaling factor for the simulation parameters that could recover the site-frequency spectra of the original model without any scaling. We then scaled the simulation parameters: mutation and recombination rates, population sizes, migration rates, and growth rates of the exponential population growth phase with a factor of 10. We did not attempt to simulate whole genomes, but 400 sub-simulations of fragments of size 10Kbp as was done for the other models. For all models described above, we sampled individuals at the end of the simulation. For the AO model, we sampled at the end by pooling the three populations: Africans, Europeans, and East Asians and taking a sample of individuals from this new population, but we also took samples from each of the three subpopulations before they were pooled.

### 2*Ns* Densities

For distributions of the population selection coefficient we considered a normal distribution, as well as inverted lognormal and gamma distributions, i.e. extending to −∞ rather than +∞. Rather than have the upper limit at zero, which would not allow for the inclusion of strictly neutral mutations, we set the upper limit at 1, so as to include the possibility of weakly advantageous mutations as well as neutral mutations. These densities are then:

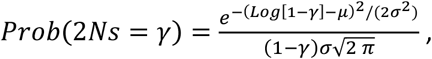

for an inverted lognormal density with maximum 1, expectation *μ* and standard deviation *σ* for the natural logarithm of 1 − *γ*, and

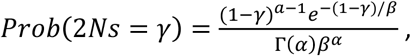

for an inverted gamma density with maximum 1, shape parameter *α* and scale parameter *β*.

We also considered a normal (gaussian) distribution. as well as a mixed distributions, that included a continuous distribution (lognormal, gamma or normal, as described) and a point mass at a value between -10 and 10. For any continuous density *f* (*γ*), the corresponding mixture with a point mass value and location given by *p*^+^ and *γ*^+^, respectively, is

*Prob* (2*Ns* = *γ*) == (1 − *p*^+^)*f*(*γ*) + *p*^+^*δ*(*γ* − *γ*^+^), where *δ*() is the Dirac delta function.

### Drosophila Applications

We extracted synonymous, non-synonymous, and short intron site-frequency spectrums from whole-genome sequencing data sets of two *Drosophila melanogaster* populations for the four autosomal chromosome arms. The North Carolina population has 200 inbred lines (39) while the Zambia collection is based on 197 haploid embryos (27). Because both data sets have missing data in some lines at some positions, allele counts at all SNPs were down sampled to the expected SFS with a uniform sample size of 160. Because of uncertainty as to which allele is truly ancestral for a given SNP, we used folded SFSs (8).

All sequence data were downloaded from the Drosophila Genome Nexus (DGN, https://www.johnpool.net/genomes.html). For each population a VCF file was constructed by first running the ‘masking package’ (available at DGN) to mask identical-by-descent or admixture tracks when present in one or many genomes, followed by the ‘snp-site’ program (40) to convert the population multi-alignment FASTA to a VCF. Because snp-site assigns the common allele as the reference, a custom script was used to assign the correct reference base. This script also ensures the genotype data conform to the standard VCF format v.4. An additional filter was applied to only keep bi-allelic SNPs genotyped on 50% or more of individuals in each population. With the DGN data in VCF format, we used GATK LiftoverVcf (41) to shift the SNP coordinate positions from *D. melanogaster* reference genome Dmel 3 to Dmel 6 (42).

We annotated SNPs with SNPEff (43). For the neutral SFS we used SNPs found in short introns (< 86 bp in length), after removing 8 bp from each side (25, 26, 44). To help ensure that the selected and neutral SNPs were as closely matched as possible for local mutational context, we followed Machado et al., (17) by pairing each candidate selective SNP with the nearest short intron (SI) SNP that had the same reference allele and flanking bases. For each candidate selected SNP, a short intron partner was identified by matching a text string that included two nucleotides from the reference genome sequence (1 bp before and after the SNP) and the SNP genotype reference/alternate allele pair (e.g. C/T). As the order of the reference and alternative allele did not matter (all analyses were carried with folded site-frequency spectra) we consider both allele pairs (e.g. C/T and T/C) together when defining the SNP mutational context. For example, if one SNP was C/T, where C was the reference allele, and T was the alternative allele (as in the VCF), and a second SNP was T/C, where T was the reference and C was the alternative, and if they both had an A nucleotide one bp before and after, both SNPs share the same mutational context (e.g. AC/TA or AT/CA). These mutational contexts were defined for every combination of SNP genotypes and flanking bases, giving a list of possible 96 mutational contexts. For each candidate selected SNP, the nearest short intron SNP with matching mutational context, and not previously sampled, was added to the short intron data. Of the three classes of sites non-synonymous, synonymous and short intron, the latter class had the fewest SNPs and was the limiting factor for the total number of SNPs included in a data set. We obtained 3100 and 6694 SI and synonymous pairs and 3242 and 7100 SI and nonsynonymous pairs for North Carolina and Zambia, respectively.

## Data and Resource Availability

The SF-Ratios program, along with simulation scripts, and other scripts used in the analysis of the site frequency spectra from the *Drosophila* populations, are available at https://github.com/jodyhey/SF_Ratios. The pipeline and scripts for making the *Drosophila* site frequency spectra are available at https://github.com/vitorpavinato/dmel_data.

## Acknowledgements

We are grateful to Adam Eyre-Walker for helpful comments. This work was supported by National Institutes of Health grant GM144468 to J. Hey.

## References

1. Lewontin RC, Krakauer J. Distribution of gene frequency as a test of the theory of the selective neutrality of polymorphisms. Genetics. 1973;74:175–95.

2. Beaumont MA, Balding DJ. Identifying adaptive genetic divergence among populations from genome scans. Mol Ecol. 2004;13(4):969–80.

3. Hudson RR, Kreitman M, Aguadé M. A test of neutral molecular evolution based on nucleotide data. Genetics. 1987;116:153–9.

4. Miyata T, Yasunaga T. Molecular evolution of mRNA: a method for estimating evolutionary rates of synonymous and amino acid substitutions from homologous nucleotide sequences and its application. J Mol Evol. 1980;16(1):23–36.

5. Wright S. The distribution of gene frequencies in populations. Proc Natl Acad Sci U S A. 1937;23:307–20.

6. Kimura M. Diffusion models in population genetics. J Appl Prob. 1964;1:177–232.

7. Sawyer SA, Hartl DL. Population genetics of polymorphism and divergence. Genetics. 1992;132:1161–76.

8. Hartl DL, Moriyama EN, Sawyer SA. Selection intensity for codon bias. Genetics. 1994;138:227–34.

9. Boyko AR, Williamson SH, Indap AR, Degenhardt JD, Hernandez RD, Lohmueller KE, et al. Assessing the evolutionary impact of amino acid mutations in the human genome. PLoS Genet. 2008;4(5):e1000083.

10. Eyre-Walker A, Woolfit M, Phelps T. The distribution of fitness effects of new deleterious amino acid mutations in humans. Genetics. 2006;173(2):891–900.

11. Loewe L, Charlesworth B, Bartolome C, Noel V. Estimating selection on nonsynonymous mutations. Genetics. 2006;172(2):1079–92.

12. Eyre-Walker A, Keightley PD. The distribution of fitness effects of new mutations. Nature Reviews Genetics. 2007;8(8):610–8.

13. Williamson SH, Hernandez R, Fledel-Alon A, Zhu L, Nielsen R, Bustamante CD. Simultaneous inference of selection and population growth from patterns of variation in the human genome. Proceedings of the National Academy of Sciences. 2005;102(22):7882–7.

14. Caicedo AL, Williamson SH, Hernandez RD, Boyko A, Fledel-Alon A, York TL, et al. Genome-wide patterns of nucleotide polymorphism in domesticated rice. PLoS Genetics. 2007;3(9):e163.

15. Bhaskar A, Wang YR, Song YS. Efficient inference of population size histories and locus-specific mutation rates from large-sample genomic variation data. Genome Research. 2015;25(2):268–79.

16. Wakeley J. Polymorphism and divergence for island-model species. Genetics. 2003;163(1):411–20.

17. Machado HE, Lawrie DS, Petrov DA. Pervasive Strong Selection at the Level of Codon Usage Bias in Drosophila melanogaster. Genetics. 2020;214(2):511–28.

18. Sendrowski J, Bataillon T. fastDFE: fast and flexible inference of the distribution of fitness effects. Mol Biol Evol. 2024;41(5):msae070.

19. Tataru P, Mollion M, Glémin S, Bataillon T. Inference of distribution of fitness effects and proportion of adaptive substitutions from polymorphism data. Genetics. 2017;207(3):1103–19.

20. Galtier N. Adaptive Protein Evolution in Animals and the Effective Population Size Hypothesis. PLoS Genetics. 2016;12(1):e1005774.

21. Eyre-Walker A, Keightley PD. Estimating the rate of adaptive molecular evolution in the presence of slightly deleterious mutations and population size change. Mol Biol Evol. 2009;26(9):2097–108.

22. Griffin TF. Distribution of the ratio of two poisson random variables: Texas Tech University; 1992.

23. Díaz-Francés E, Rubio FJ. On the existence of a normal approximation to the distribution of the ratio of two independent normal random variables. Statistical Papers. 2013;54:309–23.

24. Gravel S, Henn BM, Gutenkunst RN, Indap AR, Marth GT, Clark AG, et al. Demographic history and rare allele sharing among human populations. Proceedings of the National Academy of Sciences. 2011;108(29):11983–8.

25. Parsch J, Novozhilov S, Saminadin-Peter SS, Wong KM, Andolfatto P. On the utility of short intron sequences as a reference for the detection of positive and negative selection in Drosophila. Mol Biol Evol. 2010;27(6):1226–34.

26. Clemente F, Vogl C. Unconstrained evolution in short introns?–An analysis of genome-wide polymorphism and divergence data from Drosophila. J Evol Biol. 2012;25(10):1975–90.

27. Lack JB, Cardeno CM, Crepeau MW, Taylor W, Corbett-Detig RB, Stevens KA, et al. The Drosophila genome nexus: a population genomic resource of 623 Drosophila melanogaster genomes, including 197 from a single ancestral range population. Genetics. 2015;199(4):1229–41.

28. Lack JB, Lange JD, Tang AD, Corbett-Detig RB, Pool JE. A thousand fly genomes: an expanded Drosophila genome nexus. Mol Biol Evol. 2016;33(12):3308–13.

29. Nei M, Gojobori T. Simple methods for estimating the numbers of synonymous and nonsynonymous nucleotide substitutions. Mol Biol Evol. 1986;3:418–26.

30. Keightley PD, Eyre-Walker A. Joint inference of the distribution of fitness effects of deleterious mutations and population demography based on nucleotide polymorphism frequencies. Genetics. 2007;177(4):2251–61.

31. Zeng K, Charlesworth B. Estimating selection intensity on synonymous codon usage in a nonequilibrium population. Genetics. 2009;183(2):651–62.

32. Akaike H. A new look at the statistical model identification. IEEE Transactions on Automatic Control. 1974;19(6):716–23.

33. Duret L, Galtier N. Biased gene conversion and the evolution of mammalian genomic landscapes. Annual review of genomics and human genetics. 2009;10:285–311.

34. Huber CD, Kim BY, Marsden CD, Lohmueller KE. Determining the factors driving selective effects of new nonsynonymous mutations. Proceedings of the National Academy of Sciences. 2017;114(17):4465–70.

35. ? Ragsdale AP, Coffman AJ, Hsieh P, Struck TJ, Gutenkunst RN. Triallelic population genomics for inferring correlated fitness effects of same site nonsynonymous mutations. Genetics. 2016;203(1):513–23.

36. Kousathanas A, Keightley PD. A comparison of models to infer the distribution of fitness effects of new mutations. Genetics. 2013;193(4):1197–208.

37. Haller BC, Messer PW. SLiM 3: Forward Genetic Simulations Beyond the Wright–Fisher Model. Mol Biol Evol. 2019;36(3):632–7.

38. Wright S. Evolution in Mendelian populations. Genetics. 1931;16:97–159.

39. Mackay TFC, Richards S, Stone EA, Barbadilla A, Ayroles JF, Zhu D, et al. The Drosophila melanogaster Genetic Reference Panel. Nature. 2012;482(7384):173–8.

40. Page AJ, Taylor B, Delaney AJ, Soares J, Seemann T, Keane JA, et al. SNP-sites: rapid efficient extraction of SNPs from multi-FASTA alignments. Microbial genomics. 2016;2(4):e000056.

41. Van der Auwera GA, O’Connor BD. Genomics in the cloud: using Docker, GATK, and WDL in Terra: O’Reilly Media; 2020.

42. dos Santos G, Schroeder AJ, Goodman JL, Strelets VB, Crosby MA, Thurmond J, et al. FlyBase: introduction of the Drosophila melanogaster Release 6 reference genome assembly and large-scale migration of genome annotations. Nucleic Acids Res. 2015;43(D1):D690–D7.

43. Cingolani P, Platts A, Wang LL, Coon M, Nguyen T, Wang L, et al. A program for annotating and predicting the effects of single nucleotide polymorphisms, SnpEff: SNPs in the genome of Drosophila melanogaster strain w1118; iso-2; iso-3. Fly. 2012;6(2):80–92.

44. Halligan DL, Keightley PD. Ubiquitous selective constraints in the Drosophila genome revealed by a genome-wide interspecies comparison. Genome Research. 2006;16(7):875–84.

